## Supplementary figures and images for "Immunopeptidomics-based design of highly effective mRNA vaccine formulations against *Listeria monocytogenes*"

### Supplementary Figure 1

#### Supplementary Figure 1

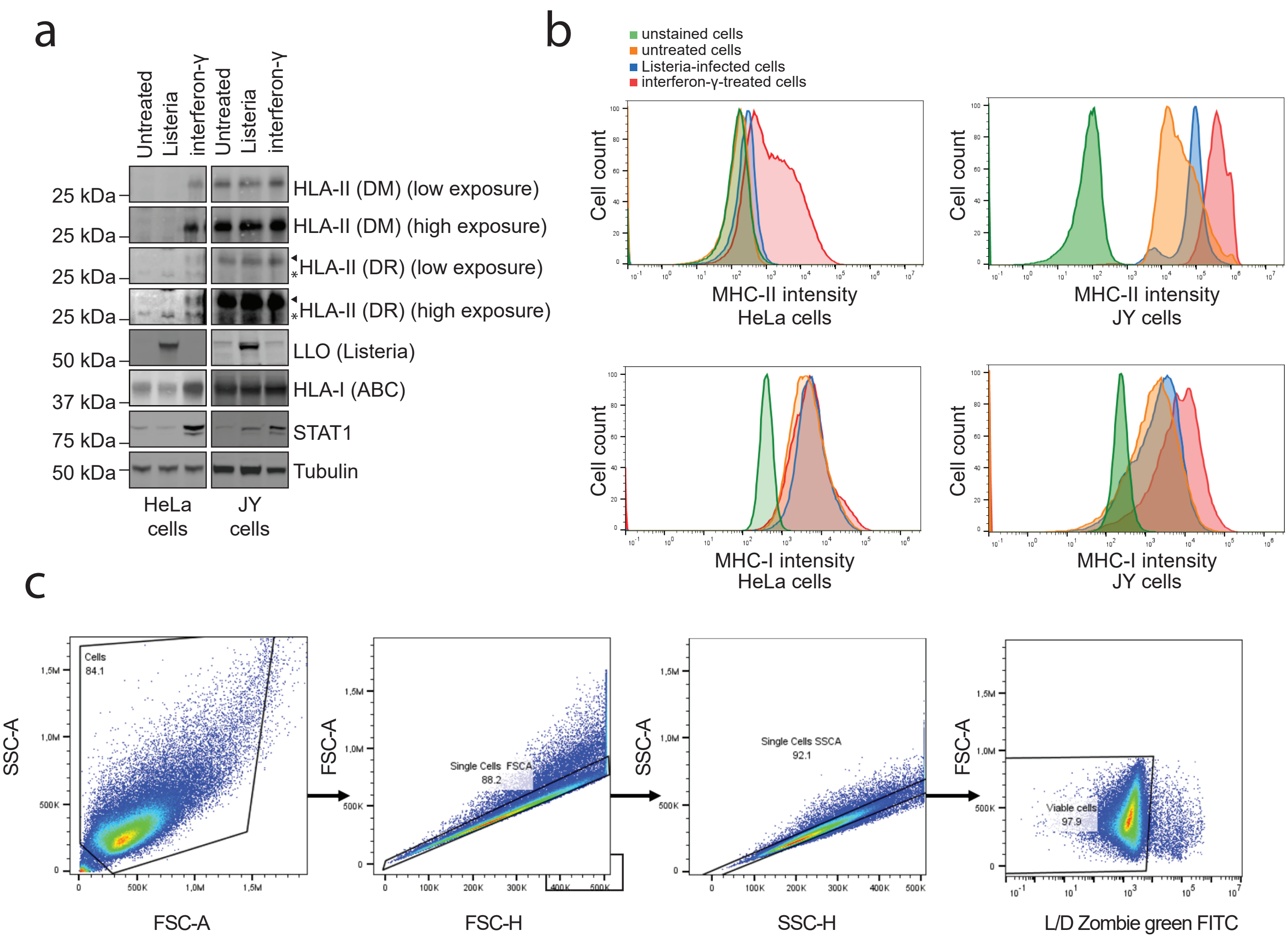

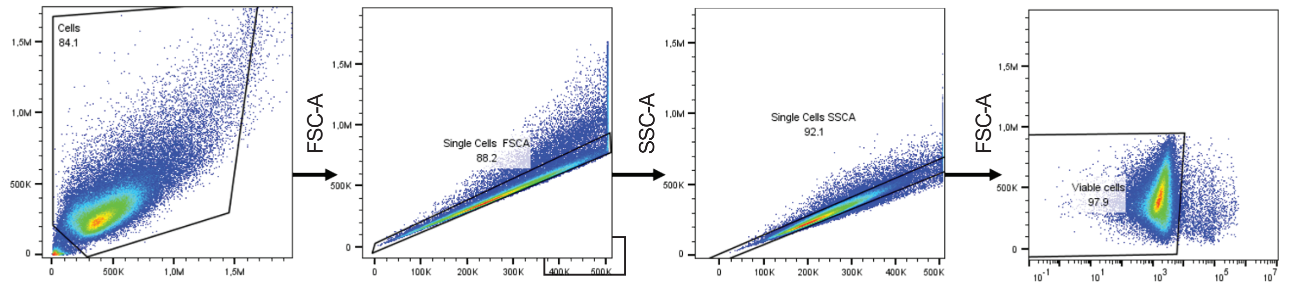

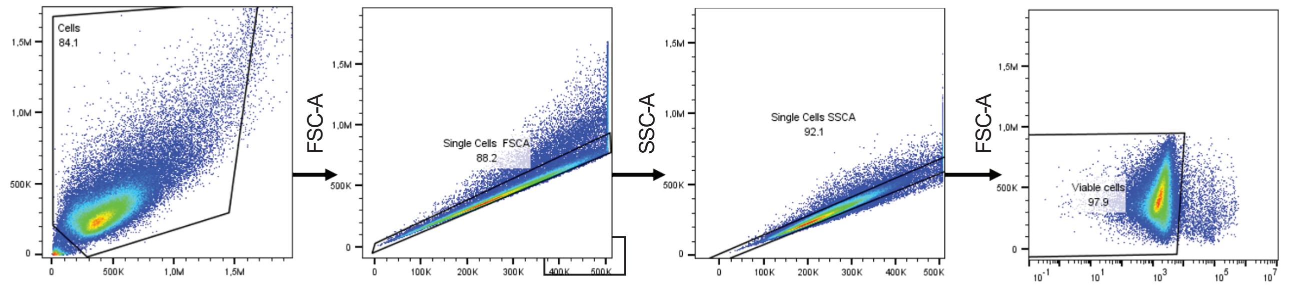

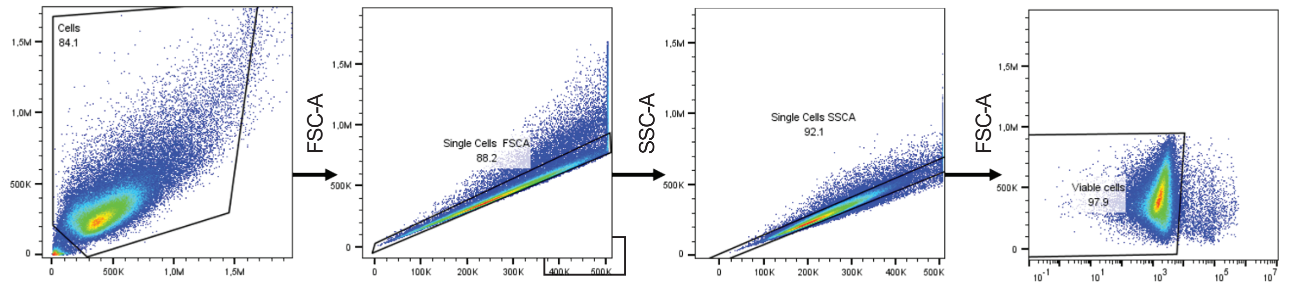

### Supplementary Figure 2

#### Supplementary Figure 2

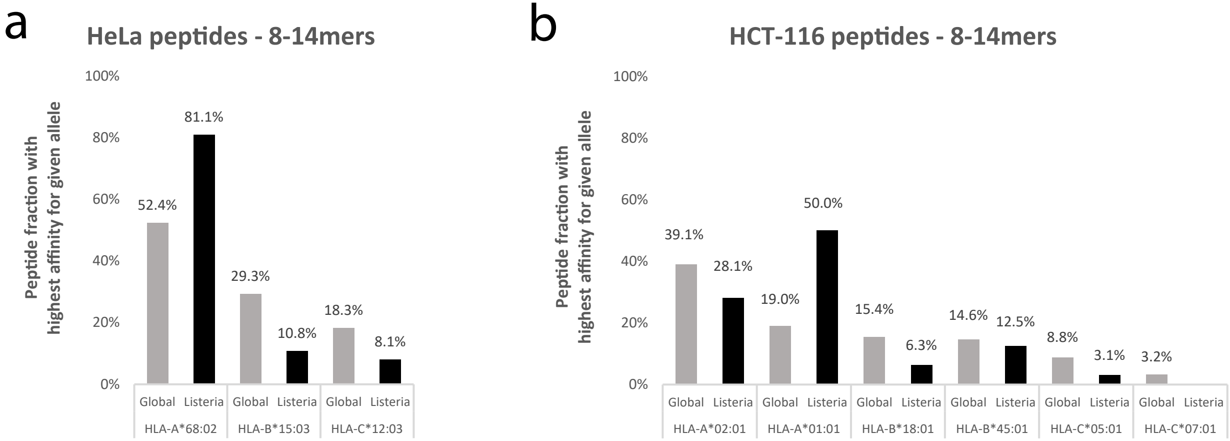

### Supplementary Figure 3

#### Supplementary Figure 3

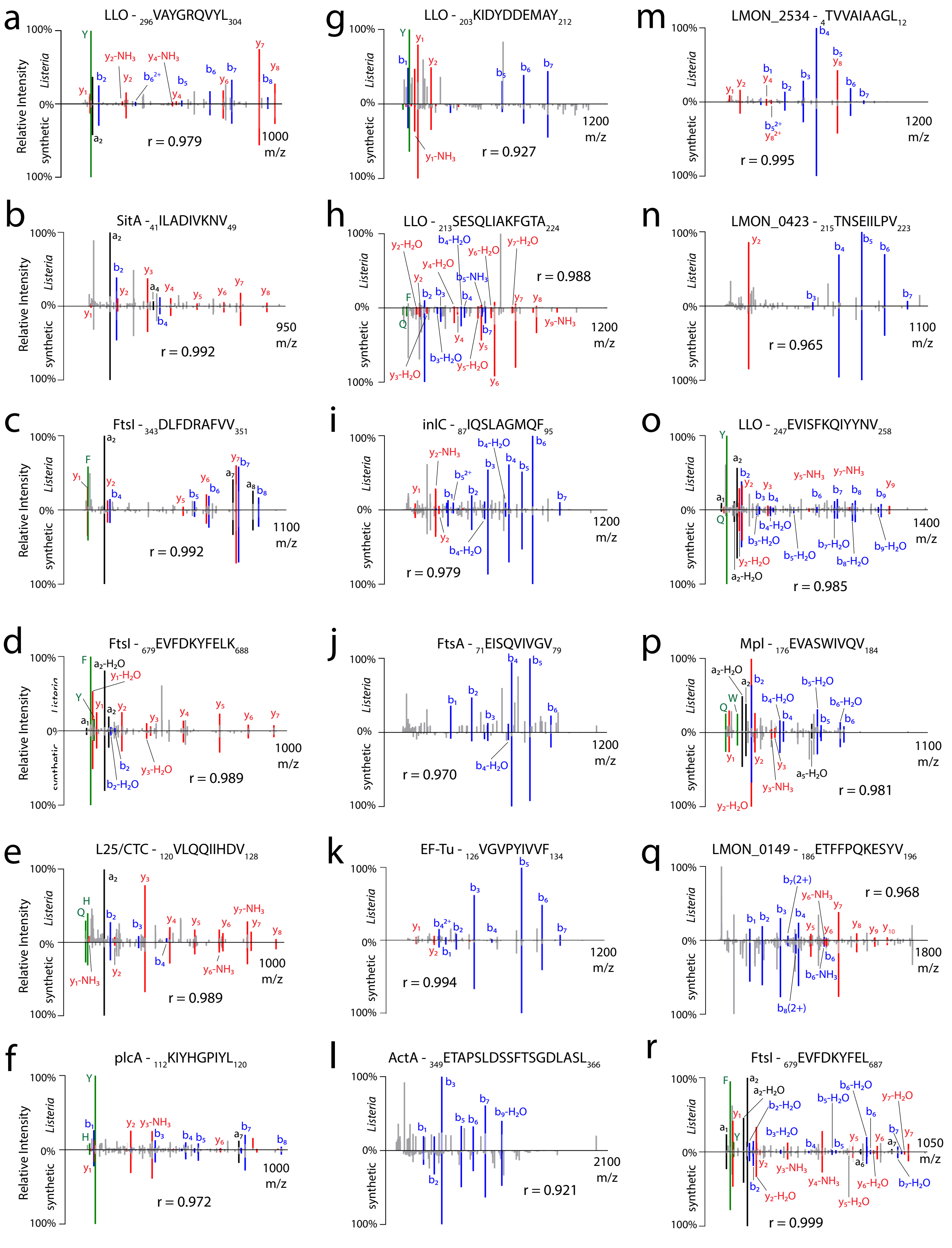

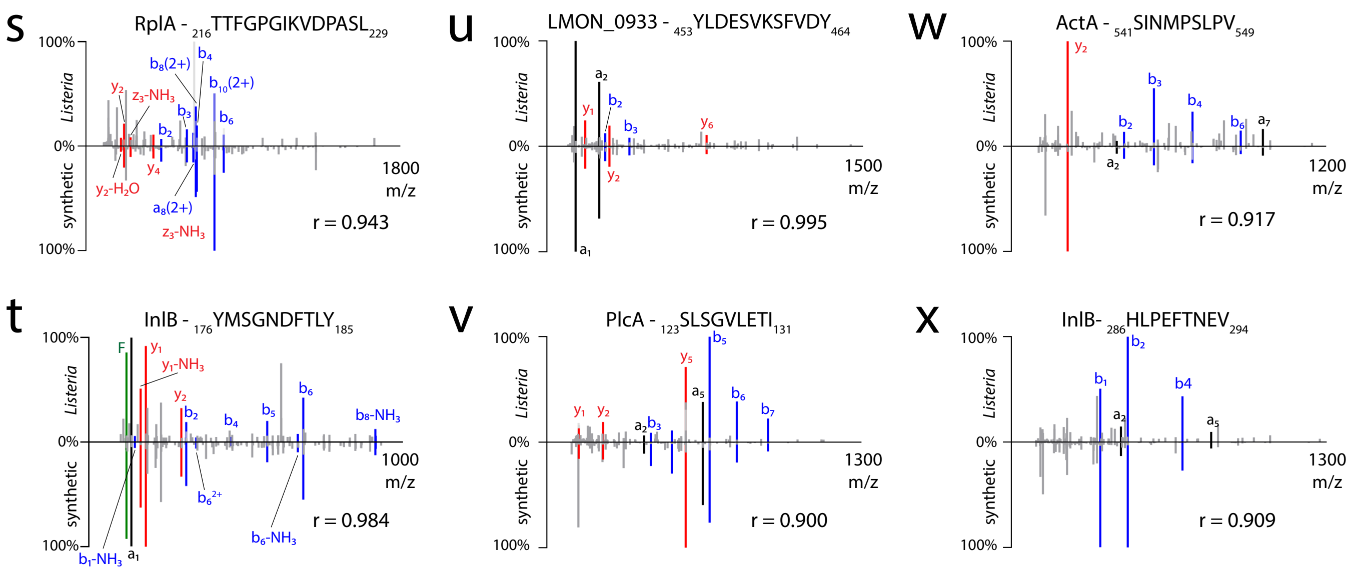

### Supplementary Figure 4

#### Supplementary Figure 4

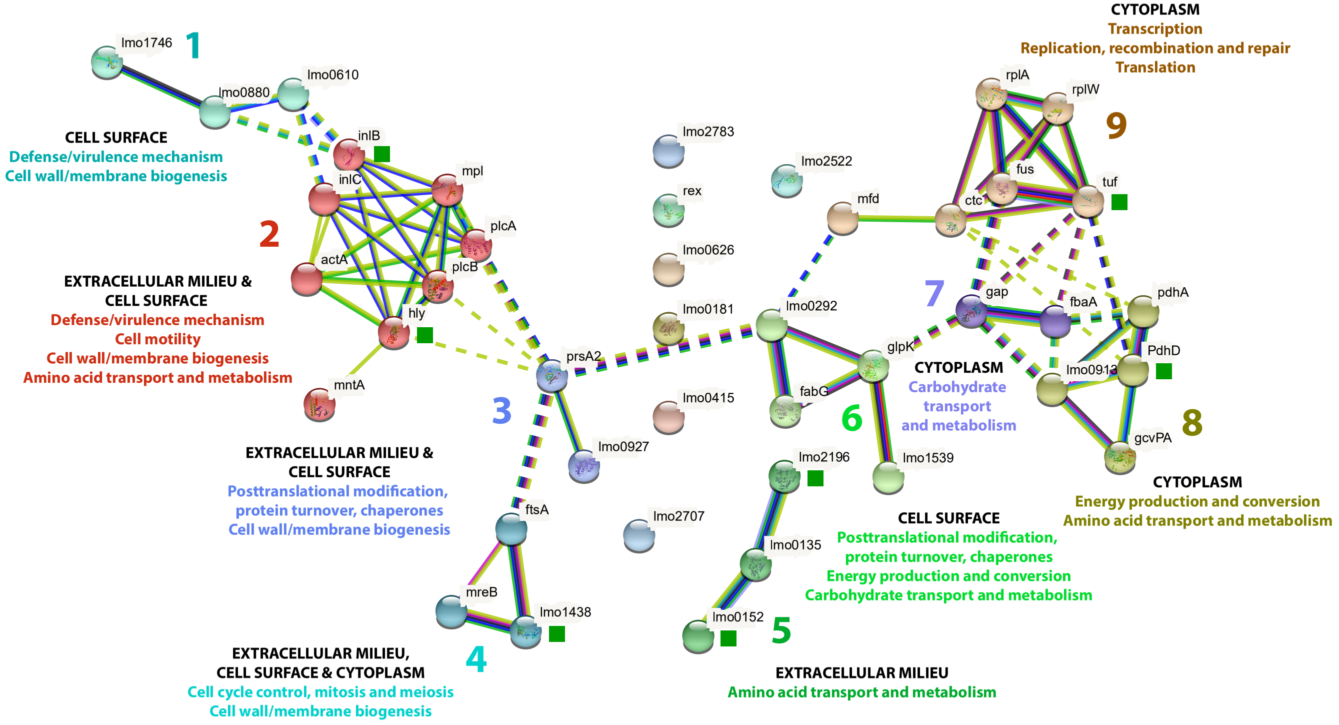

### Supplementary Figure 5

#### Supplementary Figure 5

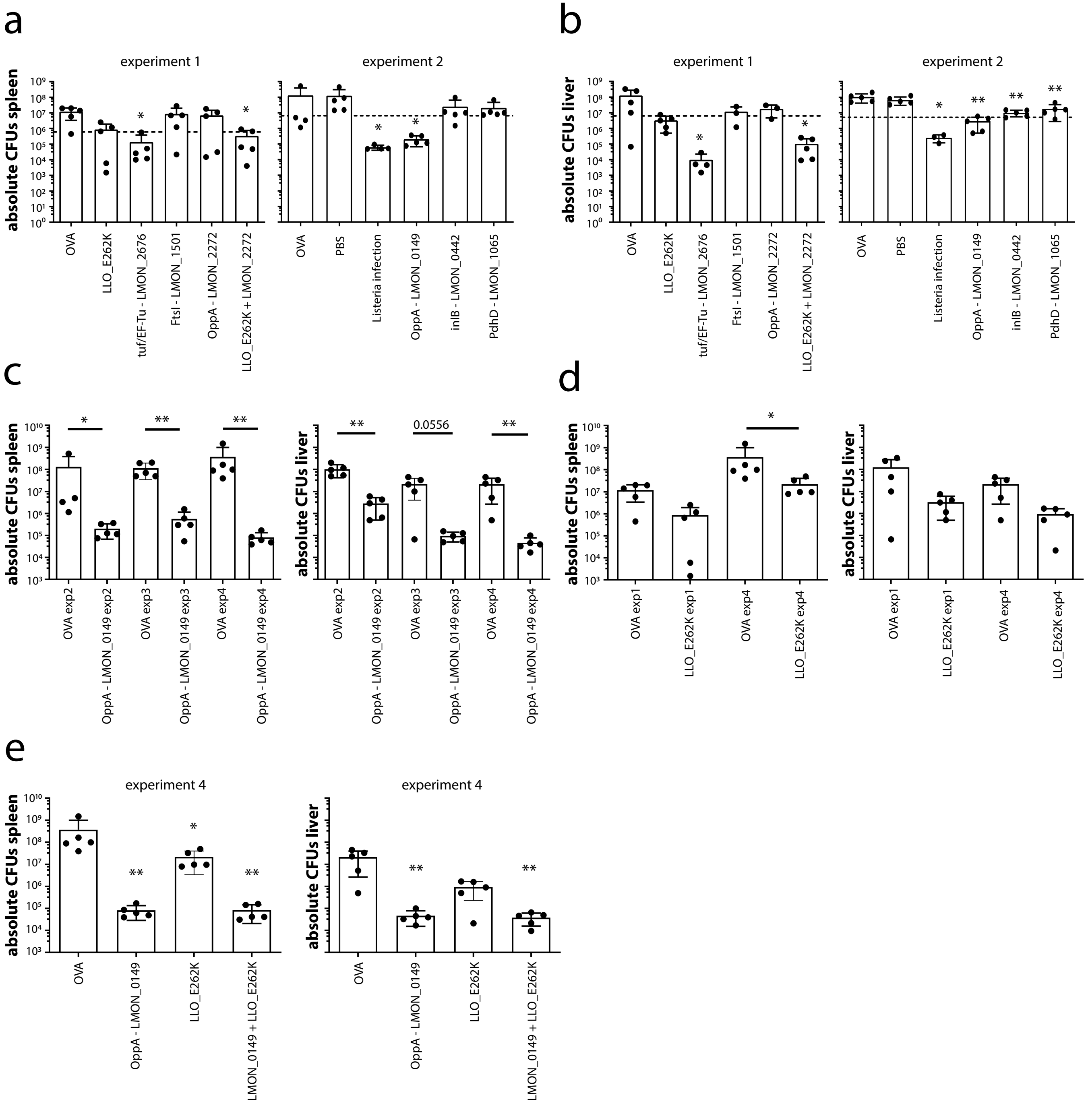

### Supplementary Figure 6

#### Supplementary Figure 6

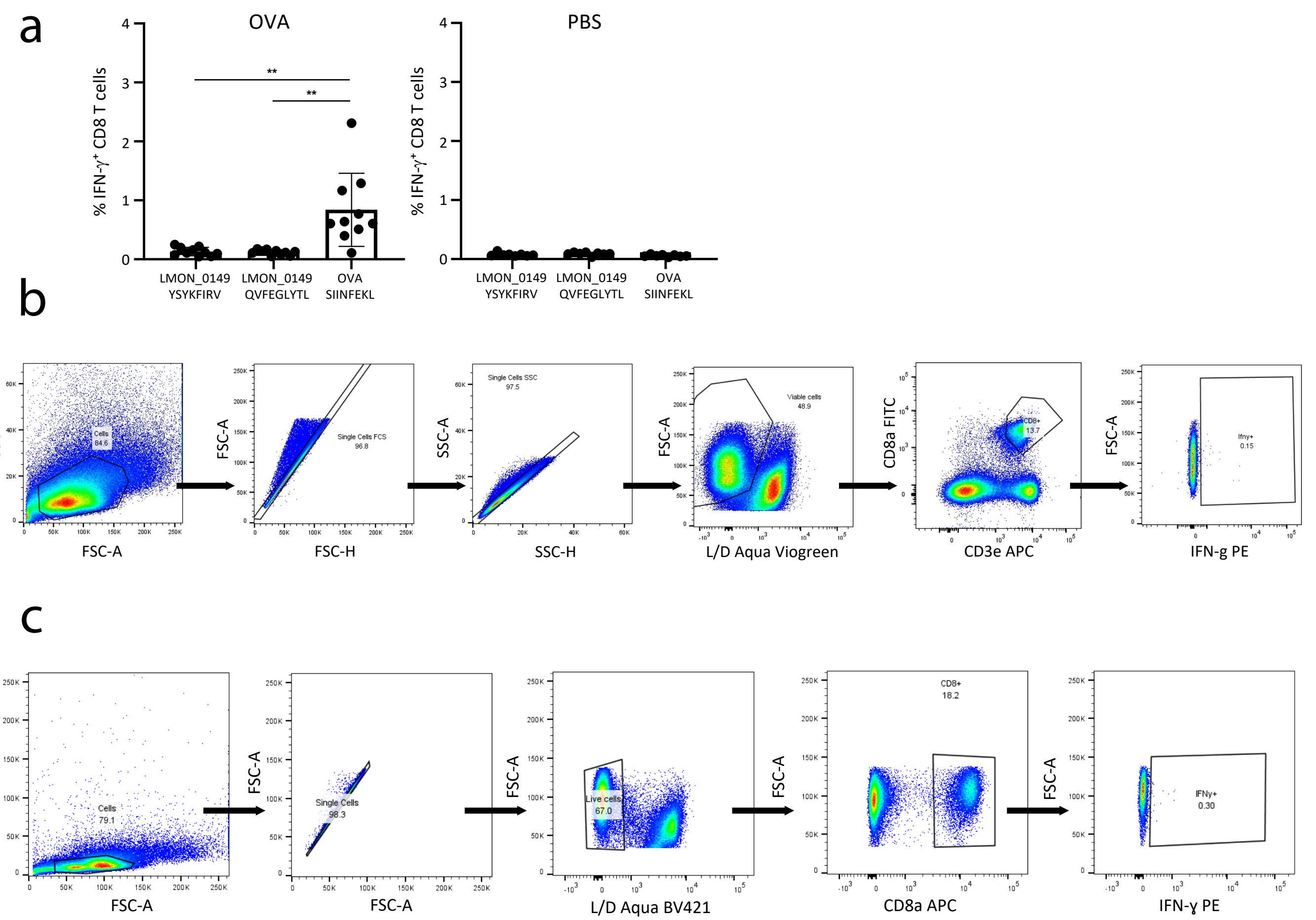

### Supplementary Figure 7

#### Supplementary Figure 7

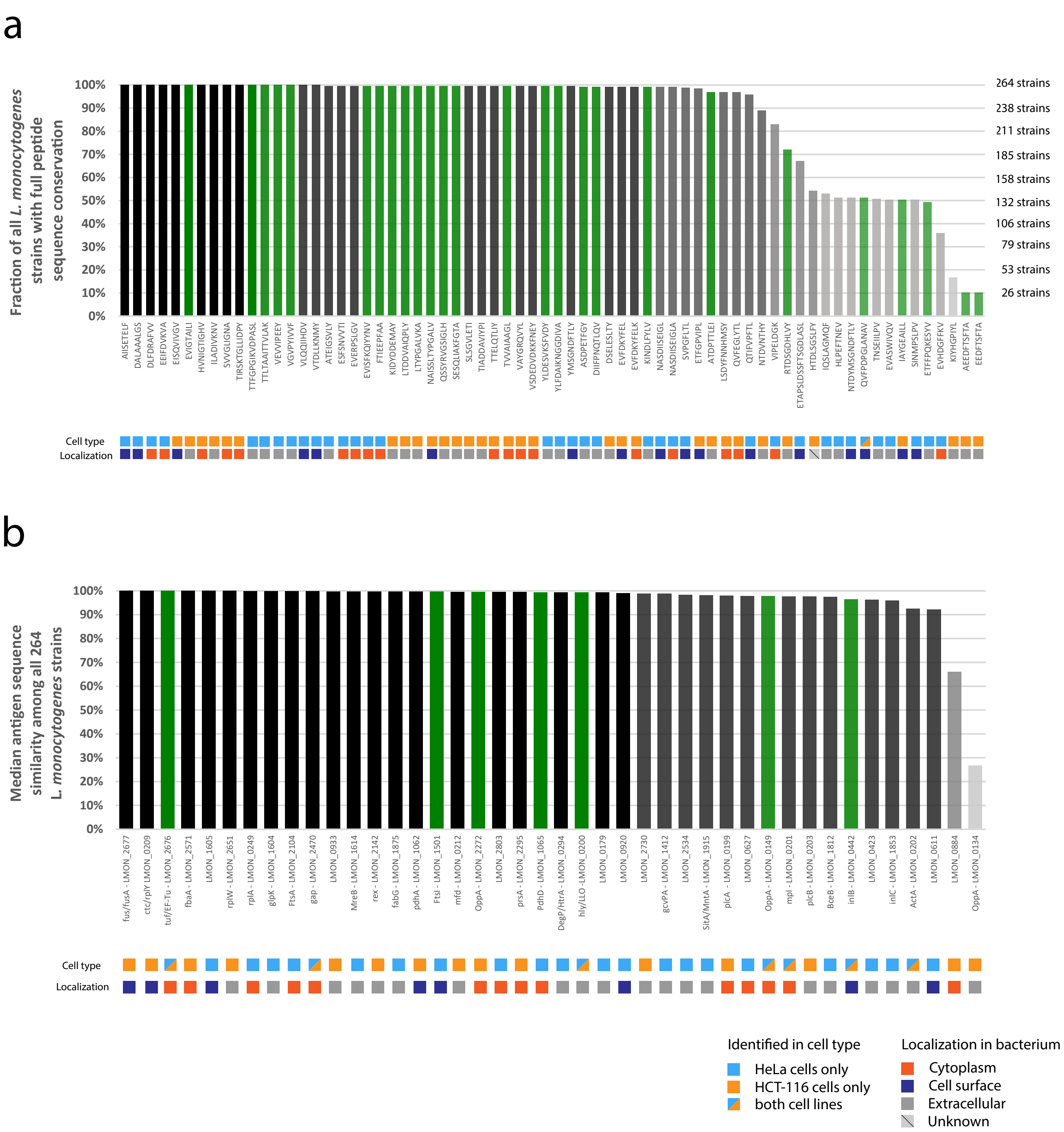
